# YamOmics: A comprehensive data resource on yam multi-omics

**DOI:** 10.1101/2024.01.23.576833

**Authors:** Jinding Liu, Mengda Wang, Yi Zhao, Danyu Shen, Qingxiang Yang, Tiegang Yang, Jianmei Yin, Longfei He, Daolong Dou

## Abstract

Yams (Dioscorea spp.) are a highly important class of horticultural crops, serving as a staple food for millions of people in Africa and contributing significantly to food security. They are also widely cultivated in East Asia as medicinal herbs, bringing substantial economic incomes. Diverse omics data play a pivotal role in advancing yam research and breeding. However, these data are often scattered, lacking in systematic organization and analysis, which underscores the need for centralized and comprehensive data management. In view of this, we gathered extensive omics data and developed the Yam Omics Database (YamOmics; https://biotec.njau.edu.cn/yamdb). The database currently offers a vast and diverse range of omics data, covering genomic, transcriptomic and plastomic data from 41 distinct yam species, along with detailed records of genomic variants from ∼1000 germplasms, and gene expression profiles from ∼200 samples. Additionally, the database features thorough annotations, encompassing aspects like genome synteny, ortholog groups, signaling pathways, gene families and protein interactions. To support yam basic biology and breeding research, it is also equipped with a suite of user-friendly online tools, including PCR primer design, CRISPR design, expression analysis, enrichment analysis, and kinship analysis tools.

## Introduction

Yams (*Dioscorea* spp.) are important horticulture crops that belong to monocotyledonous plants, but have some characteristics of dicotyledonous plants, making them important materials for fundamental botany research. Among more than 600 species of the genus *Dioscorea*, some have been domesticated for starch production, such as *D. alata, D. dumetorum, D. rotundata*, and more [1]. In some African countries, yams serve staple food, feeding millions of people, and they are also important sources of dietary diversification and are gradually increasing their market share in the global and local market (Fig. 1). On the other hand, since diosgenin saponins were first isolated from the rhizomes of *D. tokoro* in the 1930s [2], some *Dioscorea* species are also used as a favored source of medicinal plants, such as *D. nipponica, D. polystachya*, and *D. zingiberensis* [1, 3]. Over the past twenty years, worldwide production of yams has seen a twofold increase. However, this growth has been primarily driven by the expansion of cultivation areas, as opposed to enhancements in productivity (FAOSTAT 2020).

**Fig. 1.**
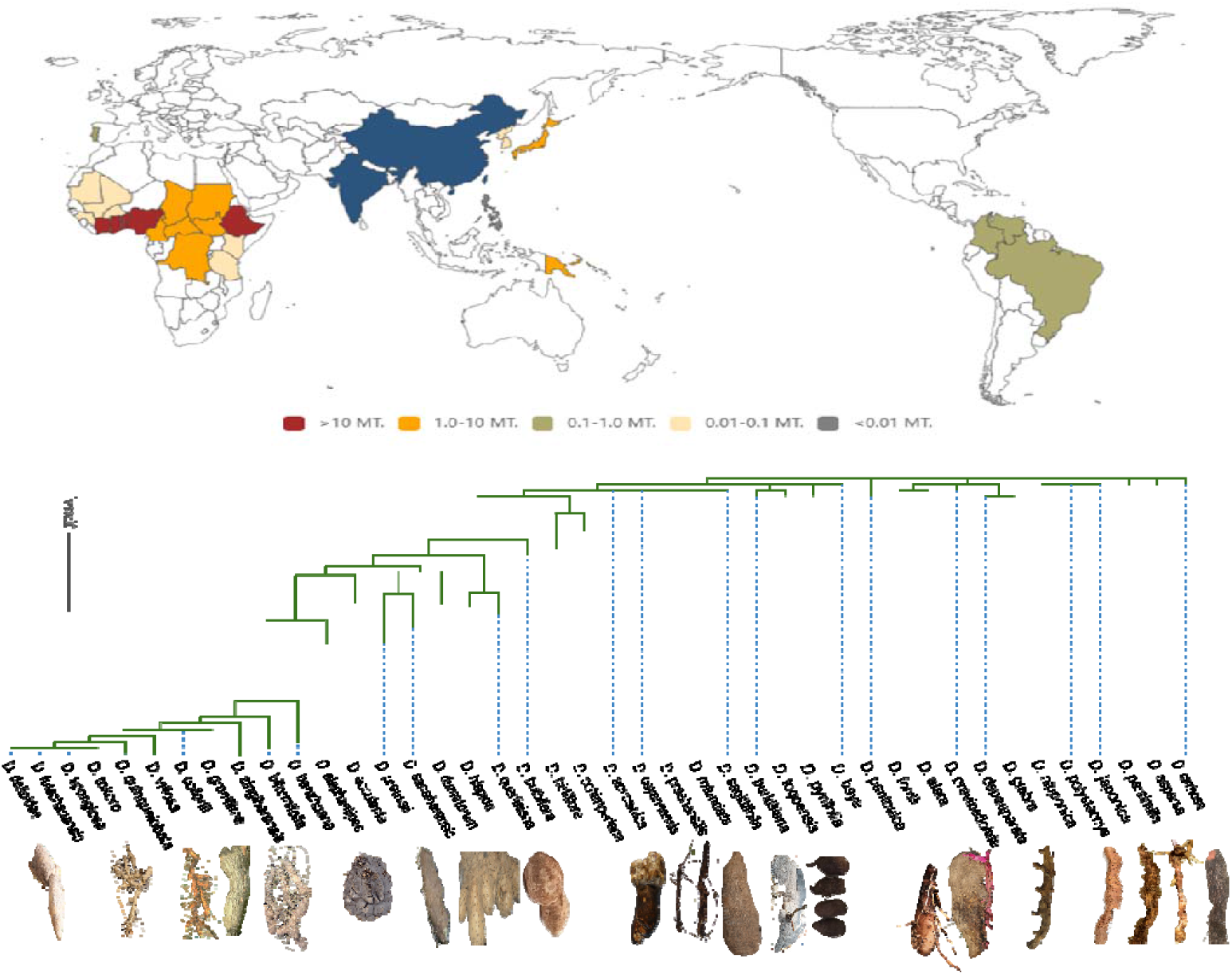
The global distribution of yam producing region, phylogeny relationship among forty-one species and some tuber phenotype.

In 2017, to facilitate genetic research and molecular breeding in yams, the first yam genome, *D. rotundata* (white Guinea yam), was sequenced to elucidate the mechanism of sex determination, to find a genomic region associated with female heterogametic sex determination [4]. Based on this finding, a molecular marker was developed for sex identification of Guinea yam plants at the seedling stage. Subsequently, in 2020, 336 accessions of white Guinea yam were resequenced. The research resolved its origin, revealing that it was a hybrid derived from crosses between the wild rainforest species *D. praehensilis* and the savannah adapted species *D. abyssinica* [5]. In 2022, the genome of *D. alata* was generated combined with a dense genetic map from African breeding populations. Using these genomic tools, users located the quantitative trait loci for resistance to anthracnose and several tuber quality traits, providing important references for yam breeding. In terms of medicinal Dioscorea species, the genome of *D. zingiberensis* was also sequenced to elucidate the biosynthesis mechanism of diosgenin saponins [6]. The comparative genomic analysis indicates that tandem duplication plus whole-genome duplication event, creating an important evolutionary resource for the diosgenin saponin biosynthetic pathway. In addition, the genomes of *D. tokoro* [7] and *D. dumetorum* [8] were also assembled and annotated to offer a large genetic data resource for their biology research. Apart from the nuclear genome, a number of chloroplast genome sequences were produced to identify the kinship relation among multiple Dioscorea species [9]. Although there are no available reference genomes for some important species, such as *D. nipponica* and *D. polystachya*, a significant amount of transcriptome sequencing dataset have been generated to explore gene function in various biological processes. In *D. nipponica*, RNA from different organs and methyl jasmonate treated rhizomes were sequenced to investigate dioscin biosynthesis [10]. To learn the molecular regulatory mechanism during microtuber formation of *D. polystachya*, RNAs from different development stages of microtuber were sequenced to reveal key pathways and hormone activities [11]. Despite the significant importance of omics data, there is no central and comprehensive database to store, analyze, mine and manage these large-scale datasets for yam research and breeding community.

Here we present YamOmics (Fig. 1), a web-based retrieval and analysis platform, covering genomic, transcriptomic and plastomic data from 41 distinct yam species, together with a large number of genomic variants from 935 yam accessions, and gene expression profiles from 191 yam samples. The database boasts three key features. First, it encompasses a comprehensive gene annotation, covering aspects like domain, family, pathway, and collinearity. Second, it features an array of sophisticated visualization tools that enable users to intuitively grasp the retrieval results. Finally, it is equipped with a suite of online tools designed to enhance the user experience and facilitate efficient database utilization, including Blast search, PCR primer design, CRISPR RNA design, expression analysis, enrichment analysis, and kinship analysis.

## Material and method

### Omics data collection and processing

We collected the genome assemblies and annotation data for 5 yam species including *D. alata, D. rotundata, D. tokoro, D. dumetorum*, and *D. zingiberensis*. The other four genome assemblies, except for *D. dumetorum*, were assembled to the chromosome level. The four assemblies were used for genomic collinearity analysis of pairwise combinations. The BLASTP program was used for identification of homologous genes between two species. The output of BLASTP was fed as evidence for MCScanX-2020-0-23[12] to identify two genomic collinearity blocks.

A total of 41 *Dioscorea* chloroplast genomes with complete assembly were collected from the CGIR database (https://ngdc.cncb.ac.cn/cgir). Global sequence alignment of these chloroplast genome sequences was conducted using mafft-7.429 [13]. The alignment files were input to phylip-3.696 [14] to build phylogenetic tree using the Neighbor-Joining algorithm. Finally, the phylogenetic tree was viewed using ITOL [15].

To obtain more genetic resources, we used Trinity-2.15.1 [16] to *de novo* assemble transcriptomes for 14 Dioscorea species that have no genome assembly available (Table S1). TransDecoder-2.1.0 was used to identify ORFs in the assembled transcripts with the assistance of Pfam database. The transcript with longest ORF was selected as representative of an assembled gene locus. BUSCO/5.3.0 [17] was used to estimate transcriptome assembly completeness.

As for variome, we collected ∼1000 resequencing datasets from *D. rotundata* and *D. alata* to call single nucleotide polymorphisms (SNPs) and short insertion and deletions (INDELs) (Table S2). Fastp-0.23.0 [18] was used to conduct quality control for raw WGS datasets. The clean datasets were aligned to reference genome with Bwa-0.7.17[19]. GATK/4.3.0 [20] was used to perform haplotype and genotype. We adopted “QD<2, QUAL<30, FS>60, MQ<40, MQRankSum<-12.5, ReadPosRankSum<-8” to filter SNPs, and adopted “QD<2, QUAL<50, FS>100, ReadPosRankSum < -20” to filter INDELs. Finally, *D. rotundata* has sufficient sequencing capacity and accession quantity, so biallelic variants with minor allele frequency (MAF) of 0.05 and max genotyping missing ratio of 0.3 were retained to construct the database. Due to insufficient the accession number and sequencing capacity, we adopted a more lenient criterion (mean depth = 5 and MAF = 0.05) to filter biallelic variants of *D. alata*. The final variants were annotated using snpEff [21].

### Functional annotation of gene annotation

We conducted various data mining and annotations to improve the usability of the database. First, primer-3-2.6.1 [22] was used to design PCR primers for each gene CDS sequence. Second, Interprocan-5.2.1 was used to identify functional domains in gene protein sequences. Third, TFBSTools [23] was used to detect transcription factor binding sites using the PFMs from the JASPAR [24] database. Fourth, we employed an Interolog-based approach to forecast protein interactions using all known experimentally confirmed interactions from the STRING database. Finally, all gene proteins were aligned to the Arabidopsis proteome (Araport11) and obtained corresponding gene names and family classifications.

### Online analysis

There are five online tools available for data analysis and retrieval. First, the database was equipped with the BLAST program for the sequence similarity retrieval. Second, DESeq2 and edgeR were employed to detect differentially expressed genes. The gene expression (TPM) matrix was normalized using the between-sample quantile (QNT) normalization method for coexpression analysis. The coexpression network was obtained through the calculation of the Pearson correlation coefficient between pairwise gene expression levels. Third, a customized perl script was used to extract all CRISPR-Cas9 guide RNAs in a specified genome editing region. BatMis-3.0 was employed to scan off-target regions in the whole genome. Crisproff-1.1.1 [25] was used to estimate off-targeting scores of Cas9 gRNAs. ClusterProfiler-3.16.1 [26] was used to conduct enrichment analysis. Finally, mafft-7.429 and FastTree-2.1 [27] were integrated into the database to identify kinship relationship among Dioscorea species whole chloroplast genome or DNA marker gene sequences.

### Database construction

YamOmic is a web application, and all data are stored and managed with the MySQL, a free relational database management system. On the server side, PHP, Perl and R are used for data processing. On the front-end side, HTML, CSS, JavaScript and JQuery are used to design interface. JavaScript plug-ins, including HighCharts and echarts, are employed to visualize retrieval and analysis results. JBrowse, a fast and scalable genome browser, is embedded to enable users to explore gene in a genomic context.

### Database content and usage

#### Database content

The database encompasses a comprehensive range of omics data from over 40 *Dioscorea* species, including genomic, transcriptomic, plastomic and variomic data (Fig. 2) Utilizing the 5 genome assemblies from *D. alata, D. rotundata, D. zingiberensis, D. tokor*, and *D. dumetorum*, we successfully identified a total of 6,303 collinear blocks through comparative genomic analysis between different species. Furthermore, we achieved *de novo* transcriptome assembly for 14 species, such as *D. polystachya, D. abyssinica, D. cirrhosa, D. <u>cochleariapiculata</u>, D. <u>communis</u>, D. <u>composita</u>, D. <u>japonica</u>, D. <u>nipponica</u>, D. <u>polygonoides</u>, D. <u>praehensilis</u>, D*. <u>soso,</u>*D. <u>sylvatica</u>, D*. <u>trifida, and</u>*D. <u>villosa</u>*, resulting in the identification of 1,062,791 genes. These genes, along with 149,061 genes annotated from the genome, have been divided into 80,629 orthologous groups. We also compiled 191 RNA-seq datasets, enabling detection of approximately 17.9 million expression levels for both *de novo* assembled and genome-annotated genes within these RNA-seq samples. Our prediction efforts led to the identification of around 10.4 million interolog-based protein-protein interactions. From 935 germplasm accessions of *D. alata* and *D. rotundata*, we called a total of 15.3 million SNPs and 1.5 million INDELs. Lastly, we have gathered the complete chloroplast genome assemblies of 41 Dioscorea species, which served as a basis for constructing a phylogenetic tree.

**Fig. 2.**
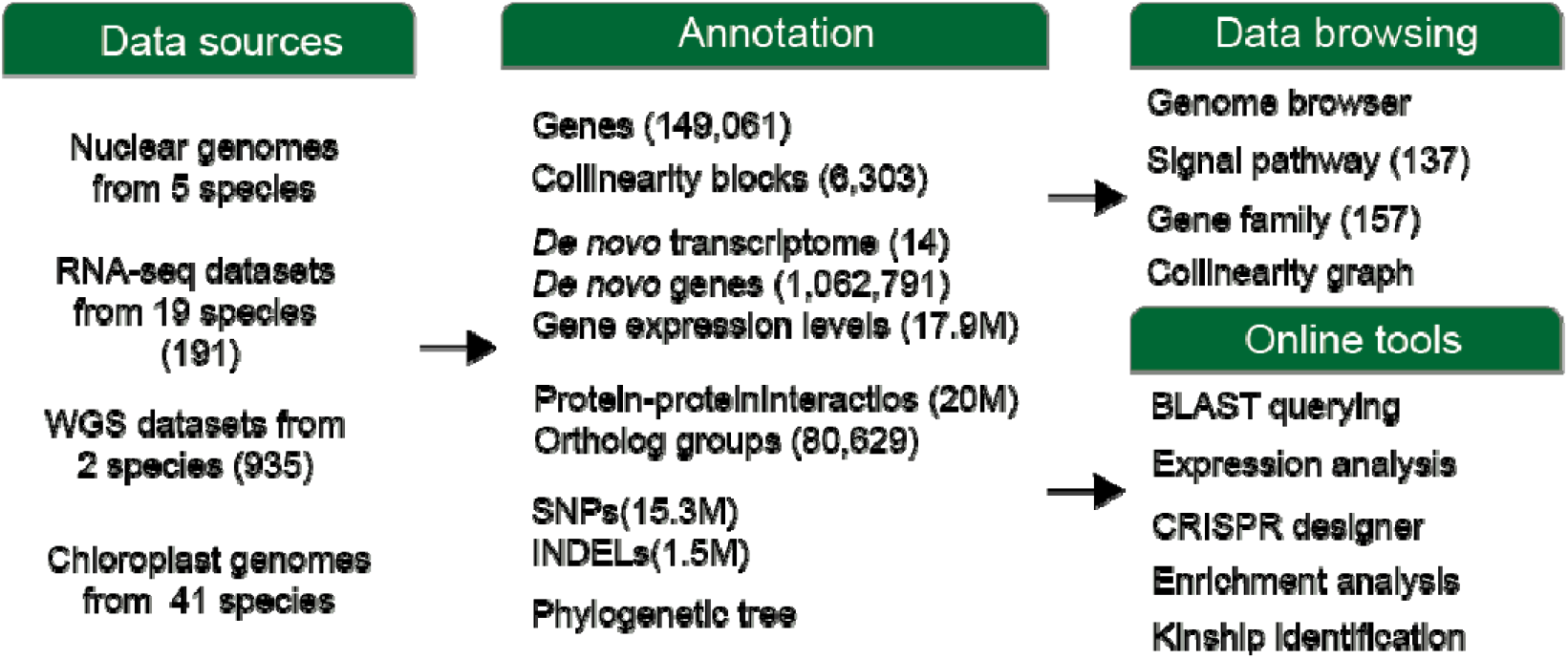
Summary of YamOmic contents and functions.

#### Database framework

YamOmic, our comprehensive database, is readily accessible at https://biotec.njau.edu.cn/yamdb. It features a user-friendly navigation menu at the top of every page. The database’s functionalities are organized into three main categories: (i) ‘Omics interfaces’ offering an immersive experience in browsing genome, transcriptome, plastome and variome data. (ii) ‘Gene interfaces’ supporting exploration of genes in the signal pathway, gene family, genome collinearity and BLAST search capabilities. (iii) ‘Analysis interfaces’ equipped with a suite of online analysis tools, such as gene differential expression/coexpression analysis, CRISPR gRNA design, gene enrichment analysis, and kinship identification within the *Dioscorea* species.

#### Omics interfaces

Through the navigation menu or function button in the home page, users can conveniently open omics page to browse related information. At the top of the genome page, there is a drop-down list box for users switching to other species. The first section displays summary information of a genome assembly, such as assembly accession number, genome size, N50, and more. Then, there a genetic linkage map showing gene density in chromosomes (Fig. 3A) and JBrowser for users to explore genes in genomic context (Fig. 3B). Next, there is a table displaying statistical information for genes in each chromosome (Fig. 3C). Clicking on the view button, user can load genes of interests into an interactive table.

**Fig. 3.**
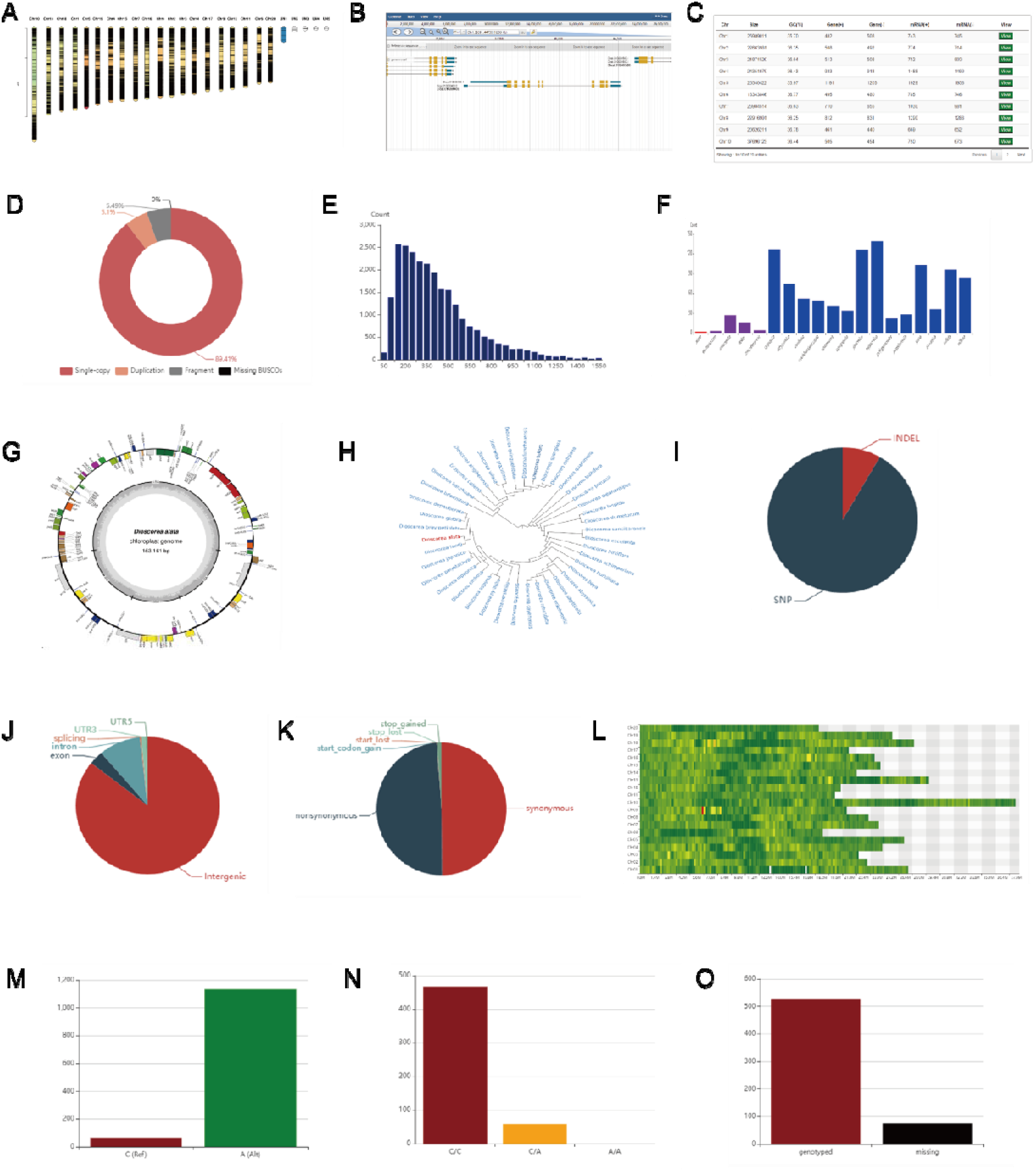
Omics data visualization. (A) Gene distribution density in chromosomes. (B) Exploration of genes in genomic context. (C) Gene table with several features. (D) Completeness estimation for *de novo* assembled transcriptome. (E) Protein sequence distribution of *de novo* assembled transcriptome. (F) Orthologous group size in genomic and transcriptomic proteomes. (G) Gene distribution in chloroplast genome. (H) Phylogenetic tree of 41 *Dioscorea* species. (I) Proportion of SNP and INDEL variants. (J) Proportion of variants in different gene feature regions. (K) Proportion of exonic variant effect. (L) Variant distribution in chromosomes. (M) Allele frequency of a variant site. (N) Genotype frequency of a variant site. (O) Missing frequency in genotyping of a variant site.

The transcriptome page has similar layout to the genome page. First, the initial section in the transcriptome page display summary information of a transcriptome assembly. Then, a pie chart (Fig. 3D) exhibits the integrity of the transcriptome assembly, and a bar chart (Fig. 3E) shows the length distribution of gene protein sequences. For *de novo* assembled transcriptomes, genes can’t be summarized by chromosomes; thus, they are categorized by orthologous groups and listed in an interactive table. There, a bar chart shows the size of an orthologous group across different species (Fig. 3F). The red bar represents the current species, the purple bars indicate species with genome available, and the blue bars indicate species with transcriptome available.

The chloroplast genome page also has a drop-down list box for users to select one of 41 species of interest. The first section is a circular genome structure diagram showing gene position (Fig. 3G). Then a circular tree diagram displays the position of a species of interest in the Dioscorea phylogenetic tree (Fig. 3F). Finally, there is an interactive table showing genes and SSRs in the chloroplast genome.

In the variome page, there are 3 pie charts showing the proportion of SNP and INDEL variants (Fig. 3I), the proportion of variants in different gene feature regions (Fig. 3J), the proportion of exonic variant effects (Fig. 3K). The following is a heatmap showing variant density across all chromosomes (Fig. 3L). Variants can be queried by genome region and the retrieved variants are listed in an interactive table. By clicking on a variant ID, users can open a page showing its details, including allele frequency (Fig. 3M), genotype frequency (Fig. 3N) and genotyping missing frequency (Fig. 3O). At the page bottom, there is a table showing the genotype of the current variant site in all germplasm accessions. In addition, the table also provides hyperlinks for users to access information of accessions and study projects.

#### Gene interfaces

To facilitate gene exploring, the collected genes can be accessed by signal pathway, gene family, genome collinearity and blast search. On the signal pathway page, there is a bar diagram showing gene distribution in 5 types of pathways including cellular processes (red), metabolism (purple), environmental information processing (blue), genetic information processing (green) and organismal systems (orange), and 18 subcategories of pathways (Fig. 4A). Through the interactive bar chart, users can quickly open a page to show a pathway of interest. In the pathway, the nodes with mapping genes are marked in red (Fig. 4B). Hovering the mouse over a node displays a message box showing gene IDs. By clicking on gene ID, users can open a page to display gene detail.

**Fig. 4.**
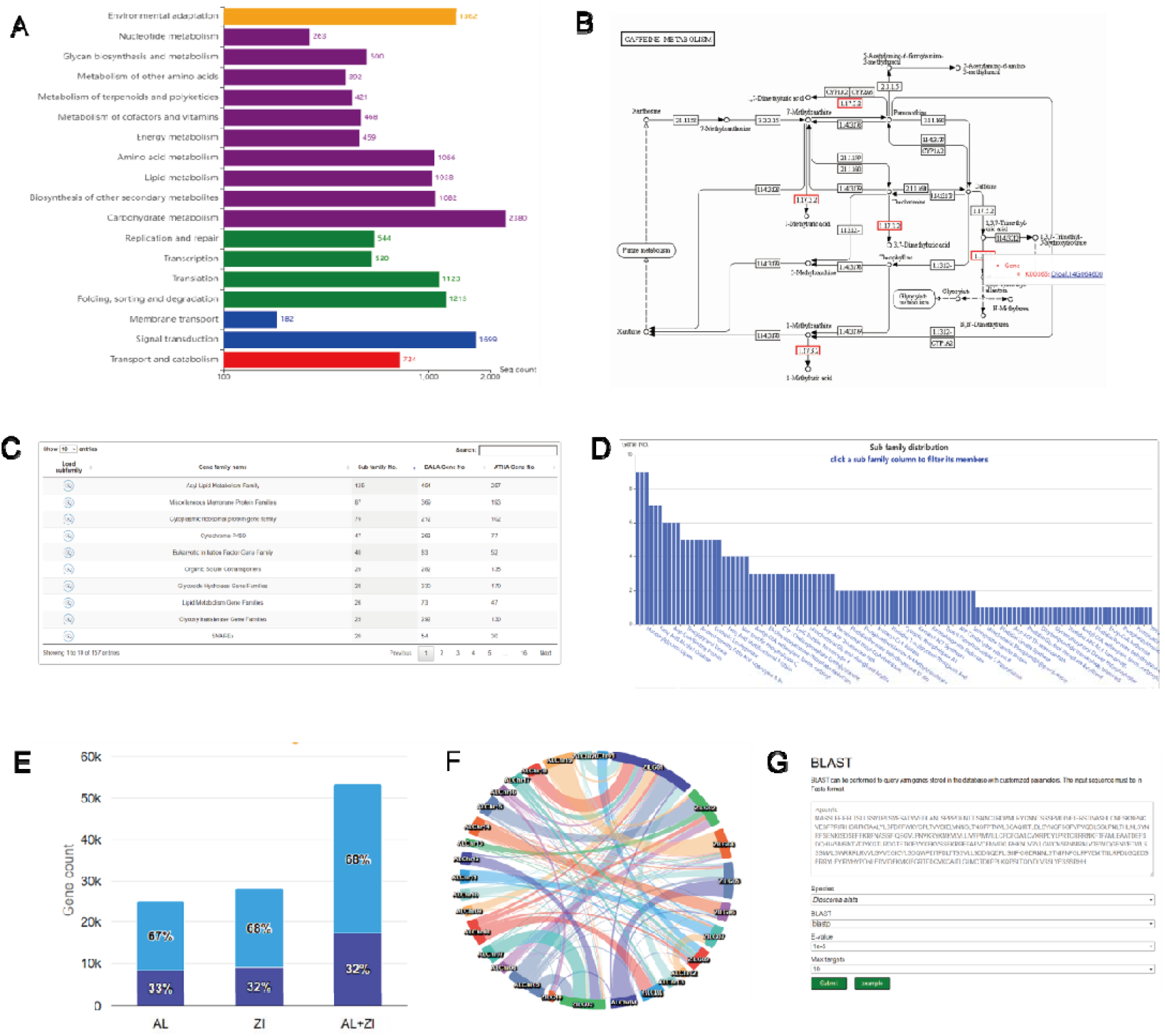
Exploring genes by multiple approaches. (A) Gene distribution in different types of signal pathways. (B) Visual chart of caffeine metabolism pathway. (C) Gene family list with family size and subfamily numbers. (D) Subfamily distribution graph of a specified family. (E) Collinear gene proportions between *Dioscorea* genomes. (F) Mapping graph of collinear blocks between Dioscorea genomes. (G) BLAST search interface.

On the gene family page, there is a table exhibiting the current species possessed gene families with family size and subfamily number (Fig. 4C). If users select a family, a bar chart appears below, showing its all subfamilies (Fig. 4D). By clicking on a subfamily bar, its gene members will be listed in a following table with hyperlinks to gene page.

In addition to pathway and gene family, genes can be explored by genome collinearity. In the genome collinearity page, it allows users to select any two genomes to list genes in their collinear blocks. In the first section of the page, an interactive bar chart is used to show the proportion of genes in collinear blocks (Fig. 4E). Then, a circle mapping graph of collinear blocks between Dioscorea genomes is present to help users to filter out genes in a specified collinear block (Fig. 4F). Clicking on a connection line, the genes in the corresponding block will be listed in a table below. Furthermore, the database is also equipped with an online BLAST program (Fig. 4G), which allows users to search homologous genes by sequence similarity.

#### Gene page

Whether starting from the omics interfaces or the gene interfaces, users can open a page to show gene details (Fig. 5). At the top of the gene page, there is a table exhibiting gene details, including gene ID, species, genome position, annotation information, and hyperlinks to download cDNA, CDS, and protein sequences (Fig. 5A). The following section include five panels named gene model, TFBS, primer, domain, expression, and interaction, respectively. The genome model panel displays the gene’s intron-exon structure (Fig. 5B). The TFBS panel offers a table to list the predicted transcription factor binding sites (Fig. 5C). The primer panel provides a table showing the recommended qPCR primers with related parameters such as temperature, stability and more (Fig. 5D). The domain panel contains a graph showing domain topology in the protein sequence and a table displaying domain details (Fig. 5E). The expression panel contains a boxplot exhibiting gene expression levels in the collected RNA-seq samples (Fig. 5F). Finally, the interaction panel also contains a bar diagram and a table exhibiting the predicted protein-protein interactions (Fig. 5G).

**Fig. 5.**
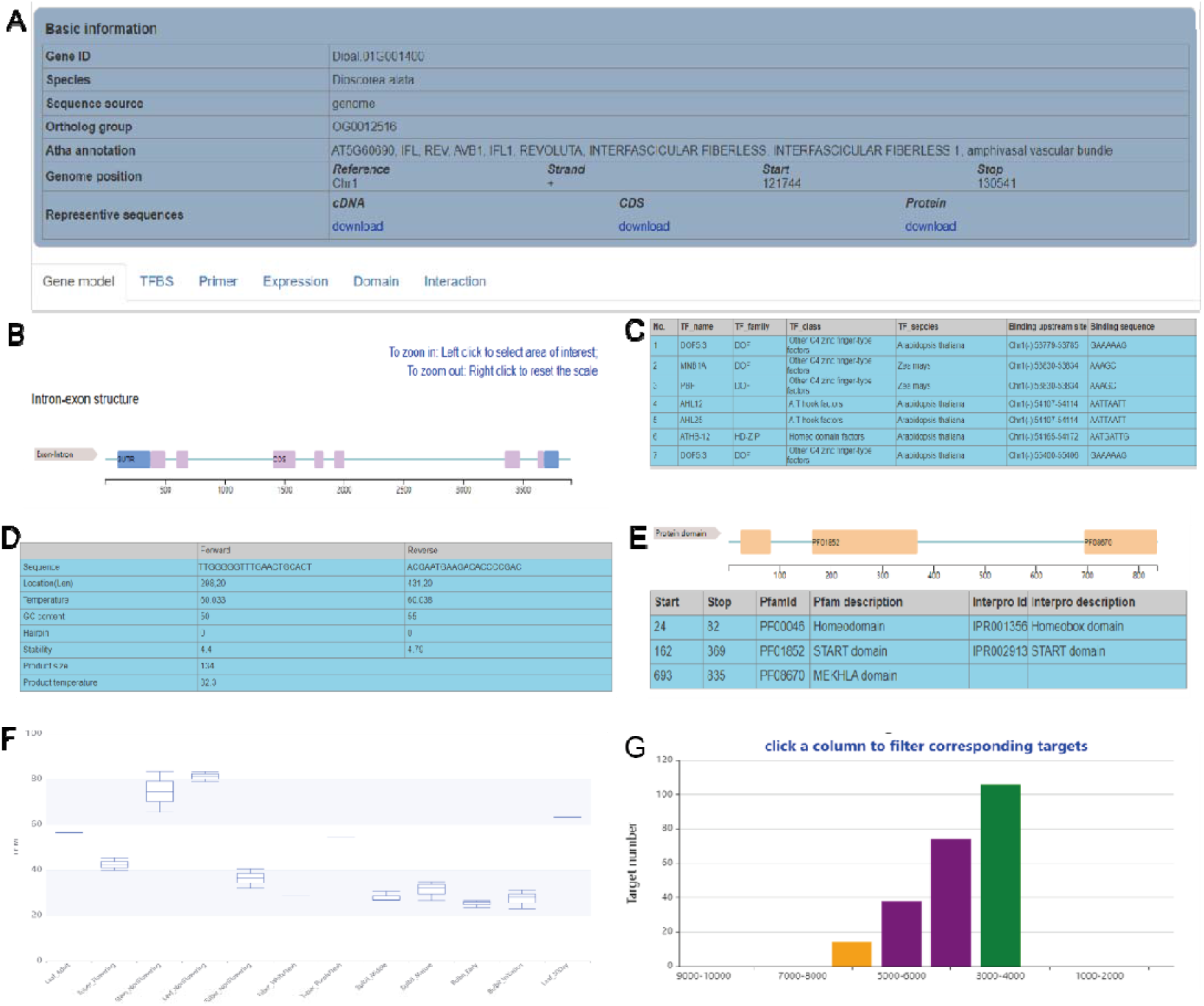
Display of gene details. (A) Summary of gene information. (B) Gene structure including UTR, CDS, and intron. (C) Transcription factor binding site table. (D) qPCR primer table. (E) Domain topology of protein sequence. (F) Boxplot showing gene expression levels. (G) Bar chart showing the distribution of predicted interaction scores.

#### Analysis interfaces

YamOmic is equipped with four online analysis tools to support yam research. In the expression analysis page, users can freely select samples and specify parameters to conduct differential gene expression and coexpression analysis (Fig. 6A). Differentially expressed genes are listed in an interactive table with log2FC, P-value, and more (Fig. 6B). By clicking on the explore button, users can further find genes that have a coexpression profile with the current gene. In addition to the coexpression genes listed in a table below (Fig. 6C), the expression profiles are visualized in a curve diagram (Fig. 6D).

**Fig. 6.**
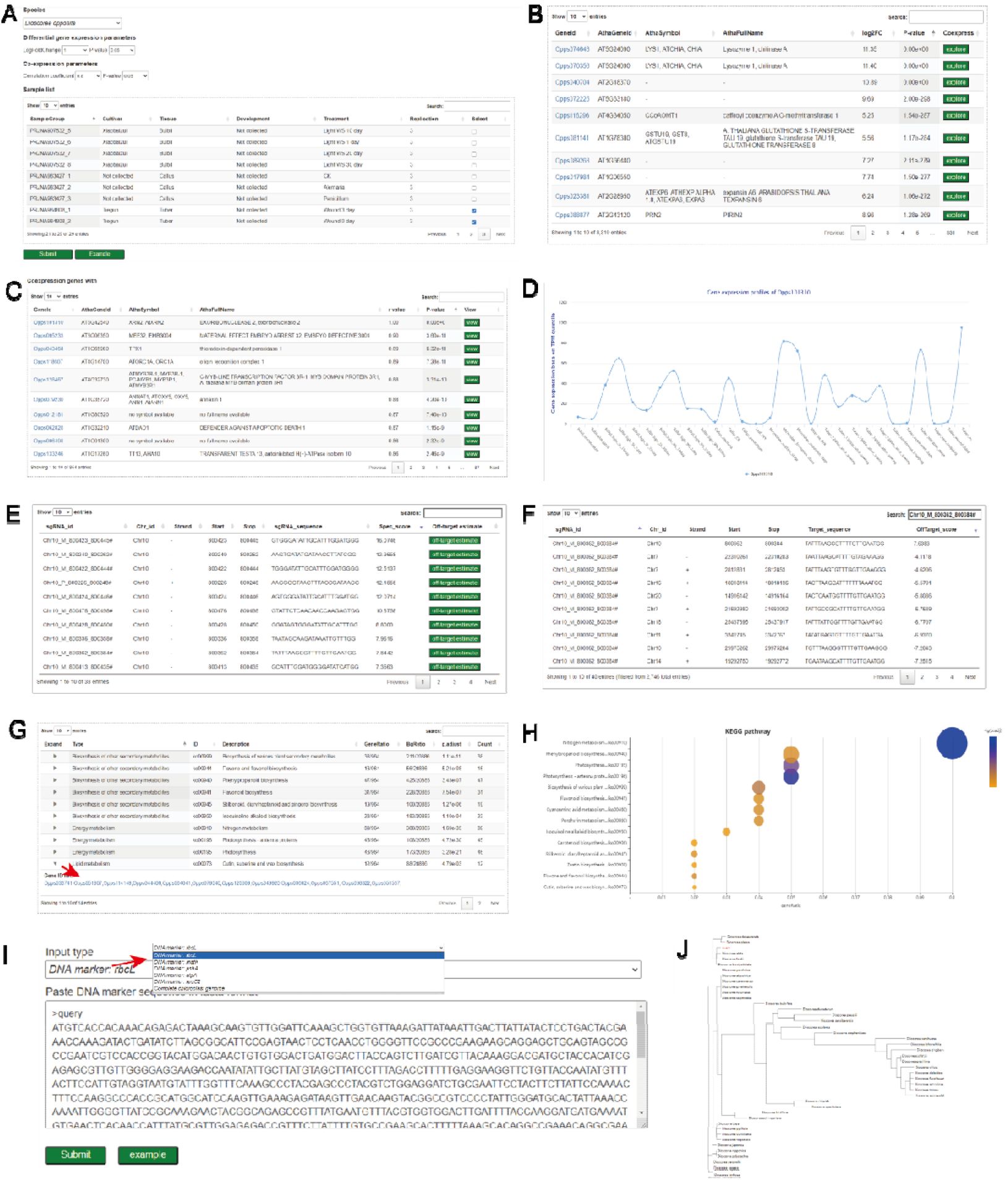
Online analysis tools. (A) Expression analysis interfaces for users to specify parameters and select samples. (B) Differentially expressed gene list. (C) Co-expression gene table. (D) Gene expression profiles in the collected RNA-seq samples. (E) CRISPR/Cas9 sgRNA table with specificity scores. (F) Table estimating off-target scores of a sgRNA. (G) Enrichment analysis results table. (H) A scatter plot to visualize enrichment analysis results. (I) Kinship identification interfaces for users to paste DNA marker sequences or upload a chloroplast genome sequence file. (J) Phylogenetic tree showing the position of a querying sequence among 41 Dioscorea species.

To help users investigate gene function, the database contains an extremely user-friendly CRISPR/Cas9 sgRNA design tool. By simply specifying a genomic region, including the chromosome number and interval coordinates, users can acquire sgRNAs tailored for editing that specific area (Fig. 6E). Additionally, the tool offers valuable insights into the editing efficiency and potential off-target effects (Fig. 6F).

Enrichment analysis is used to study the overall function of a gene set, which is commonly based on gene ontology terms or pathways. If users upload a set of gene IDs, the tool can return a table showing enrichment analysis results with significance levels and gene ratios (Fig. 6G). In addition, the scatter plot below is used to visualize the analysis results (Fig. 6H).

The kinship identification tool has been meticulously crafted to assist users in discerning the phylogenetic relationships among Dioscorea accessions. To achieve this goal, users can paste marker gene DNA sequences from the chloroplast genome including rbcL, matK, psbA, and more, or upload a complete chloroplast genome sequence file (Fig. 6I). The kinship tool can construct a phylogenetic tree online, in which the queried sequence/genome is marked in red (Fig. 6J).

## Conclusion and future directions

In summary, we have constructed the YamOmic database, which serves as a central portal for Dioscorea species and provides comprehensive genome, transcriptome, plastome, and variome resources. The rich data resource, diverse retrieval, visualization, and analysis functions make it a one-stop resource for yam researchers and breeders. As for future directions, the database will be continuously updated as more data, including phenotypic and epigenetic data, become available.

## Acknowledgements

This study was supported by grants from the China Agriculture Research System (CARS-21), the National Natural Science Foundation of China (32270208, and 32230089) and the Fundamental Research Funds for the Central Universities (KYCXJC2023001 and KYQN2023039).

## Author contributions

D.D. designed and managed the project. J.L., M.W., Y.Z. and D.S. collected and conducted data analysis. J.L., M.W., Y.Z. and D.S. constructed the database. Q.Y., T.Y., J.Y. and L.H. participated in discussions. J.L., and D.D. wrote the manuscript.

## Data Availability Statement

All data are available at: https://biotec.njau.edu.cn/yamdb.

## Conflict of interest

The authors declare that they have no conflict of interest.

## Notes

https://biotec.njau.edu.cn/yamdb

### Competing Interest Statement

The authors have declared no competing interest.

## References

1. Epping J, Laibach N. An underutilized orphan tuber crop-Chinese yam : a review. Planta 252, 58 (2020).

2. Marker RE, Tsukamoto T, DL. T C S: Diosgenin1. Journal of the American Chemical Society, 62, 2525–2532 (1940).

3. Zhai C, Lu Q, Chen X, Peng Y, Chen L, Du S: Molecularly imprinted layer-coated silica nanoparticles toward highly selective separation of active diosgenin from Dioscorea nipponica Makino. J Chromatogr A 2009, 1216(12):2254–2262.

4. Tamiru M, Natsume S, Takagi H, White B, Yaegashi H, Shimizu M, Yoshida K, Uemura A, Oikawa K, Abe A et al: Genome sequencing of the staple food crop white Guinea yam enables the development of a molecular marker for sex determination. Bmc Biol 2017, 15.

5. Sugihara Y, Darkwa K, Yaegashi H, Natsume S, Shimizu M, Abe A, Hirabuchi A, Ito K, Oikawa K, Tamiru-Oli M et al: Genome analyses reveal the hybrid origin of the staple crop white Guinea yam (). P Natl Acad Sci USA 2020, 117(50):31987–31992.

6. Li Y, Tan C, Li Z, Guo J, Li S, Chen X, Wang C, Dai X, Yang H, Song W et al: The genome of Dioscorea zingiberensis sheds light on the biosynthesis, origin and evolution of the medicinally important diosgenin saponins. Hortic Res 2022, 9:uhac165.

7. Satoshi Natsume, Hiroki Yaegashi, Yu Sugihara, Akira Abe, Motoki Shimizu, Kaori Oikawa, Benjamen White, Aoi Kudoh, Terauchi R: Whole genome sequencing of a wild yam species Dioscorea tokoro reveals a genomic region associated with sex. bioRxiv 2022.

8. Siadjeu C, Pucker B, Viehover P, Albach DC, Weisshaar B: High Contiguity De Novo Genome Sequence Assembly of Trifoliate Yam (Dioscorea dumetorum) Using Long Read Sequencing. Genes (Basel) 2020, 11(3).

9. Zhao Z, Wang X, Yu Y, Yuan S, Jiang D, Zhang Y, Zhang T, Zhong W, Yuan Q, Huang L: Complete chloroplast genome sequences of Dioscorea: Characterization, genomic resources, and phylogenetic analyses. PeerJ 2018, 6:e6032.

10. Sun W, Wang B, Yang J, Wang W, Liu A, Leng L, Xiang L, Song C, Chen S: Weighted Gene Co-expression Network Analysis of the Dioscin Rich Medicinal Plant Dioscorea nipponica. Front Plant Sci 2017, 8:789.

11. Li J, Zhao X, Dong Y, Li S, Yuan J, Li C, Zhang X, Li M: Transcriptome Analysis Reveals Key Pathways and Hormone Activities Involved in Early Microtuber Formation of Dioscorea opposita. Biomed Res Int 2020, 2020:8057929.

12. Wang Y, Tang H, Debarry JD, Tan X, Li J, Wang X, Lee TH, Jin H, Marler B, Guo H et al: MCScanX: a toolkit for detection and evolutionary analysis of gene synteny and collinearity. Nucleic Acids Res 2012, 40(7):e49.

13. Rozewicki J, Li S, Amada KM, Standley DM, Katoh K: MAFFT-DASH: integrated protein sequence and structural alignment. Nucleic Acids Res 2019, 47(W1):W5–W10.

14. Shimada MK, Nishida T: A modification of the PHYLIP program: A solution for the redundant cluster problem, and an implementation of an automatic bootstrapping on trees inferred from original data. Mol Phylogenet Evol 2017, 109:409–414.

15. Zhou T, Xu K, Zhao F, Liu W, Li L, Hua Z, Zhou X: itol.toolkit accelerates working with iTOL (Interactive Tree of Life) by an automated generation of annotation files. Bioinformatics 2023, 39(6).

16. Grabherr MG, Haas BJ, Yassour M, Levin JZ, Thompson DA, Amit I, Adiconis X, Fan L, Raychowdhury R, Zeng QD et al: Full-length transcriptome assembly from RNA-Seq data without a reference genome. Nat Biotechnol 2011, 29(7):644–U130.

17. Manni M, Berkeley MR, Seppey M, Simao FA, Zdobnov EM: BUSCO Update: Novel and Streamlined Workflows along with Broader and Deeper Phylogenetic Coverage for Scoring of Eukaryotic, Prokaryotic, and Viral Genomes. Molecular Biology and Evolution 2021, 38(10):4647–4654.

18. Chen S, Zhou Y, Chen Y, Gu J: fastp: an ultra-fast all-in-one FASTQ preprocessor. Bioinformatics 2018, 34(17):i884–i890.

19. Jung Y, Han D: BWA-MEME: BWA-MEM emulated with a machine learning approach. Bioinformatics 2022, 38(9):2404–2413.

20. De Summa S, Malerba G, Pinto R, Mori A, Mijatovic V, Tommasi S: GATK hard filtering: tunable parameters to improve variant calling for next generation sequencing targeted gene panel data. BMC Bioinformatics 2017, 18(Suppl 5):119.

21. Cingolani P, Platts A, Wang le L, Coon M, Nguyen T, Wang L, Land SJ, Lu X, Ruden DM: A program for annotating and predicting the effects of single nucleotide polymorphisms, SnpEff: SNPs in the genome of Drosophila melanogaster strain w1118; iso-2; iso-3. Fly (Austin) 2012, 6(2):80–92.

22. Koressaar T, Lepamets M, Kaplinski L, Raime K, Andreson R, Remm M: Primer3_masker: integrating masking of template sequence with primer design software. Bioinformatics 2018, 34(11):1937–1938.

23. Tan G, Lenhard B: TFBSTools: an R/bioconductor package for transcription factor binding site analysis. Bioinformatics 2016, 32(10):1555–1556.

24. Rauluseviciute I, Riudavets-Puig R, Blanc-Mathieu R, Castro-Mondragon JA, Ferenc K, Kumar V, Lemma RB, Lucas J, Cheneby J, Baranasic D et al: JASPAR 2024: 20th anniversary of the open-access database of transcription factor binding profiles. Nucleic Acids Res 2024, 52(D1):D174–D182.

25. Carlson-Stevermer J, Kelso R, Kadina A, Joshi S, Rossi N, Walker J, Stoner R, Maures T: CRISPRoff enables spatio-temporal control of CRISPR editing. Nat Commun 2020, 11(1):5041.

26. Wu T, Hu E, Xu S, Chen M, Guo P, Dai Z, Feng T, Zhou L, Tang W, Zhan L et al: clusterProfiler 4.0: A universal enrichment tool for interpreting omics data. Innovation (Camb) 2021, 2(3):100141.

27. Chu G, Warnow T: SCAMPP+FastTree: improving scalability for likelihood-based phylogenetic placement. Bioinform Adv 2023, 3(1):vbad008.

